# How Well Can AI and Physics-Based Simulations Predict the Probability a Cryptic Pocket Is Open?

**DOI:** 10.64898/2026.01.21.700870

**Authors:** Si Zhang, Justin J. Miller, Gregory R. Bowman

## Abstract

Identifying and understanding cryptic pockets remains a compelling goal in drug discovery, as they can offer new avenues for targeting proteins that are otherwise challenging to modulate. While artificial intelligence (AI) methods for structure prediction have been revolutionary, they have not learned enough physics to characterize the rest of a protein’s conformational ensemble, such as structures with cryptic pockets. This limitation has motivated the development of new AI-based methods for characterizing ensembles, or properties thereof, such as AlphaFlow, BioEmu, PocketMiner, and CryptoBank. Here, we benchmark how well these models and physics-based molecular dynamics (MD) simulations can recapitulate the thermodynamics of known cryptic pockets in Ebola VP35 and TEM β-lactamase. These two pockets were chosen because the probability that they are open and the impact of point mutations have been well characterized experimentally. Multiple methods are remarkably successful at predicting whether a mutation will increase or decrease the probability of cryptic pocket opening. However, none can reliably predict the absolute probability of pocket opening. MD is very close for the two wild-type proteins but all the methods struggle for pockets with small probabilities of opening experimentally (e.g. less than 1%). BioEmu and PocketMiner capture some trends between variants with experimental populations over 1% but have systematic errors and poorer performance for rare pockets (e.g. <1% open experimentally). These results highlight the promise of AI- and simulation-based strategies for cryptic pocket characterization while pointing to the need for further improvements to achieve robust predictors.

## Introduction

Cryptic pockets are dynamic binding sites that are usually closed but open some fraction of the time due to thermal fluctuations of a protein’s structure. They are often invisible in experimentally-derived structures, which resolve what a protein typically looks like rather than more rare states, making cryptic pockets difficult to capture experimentally^1,2^. Despite these challenges, identifying such elusive pockets can create new opportunities for drug discovery by revealing binding sites on proteins historically deemed “undruggable”, providing a means to achieve specificity, or enabling allosteric regulation of functional regions^3–11^.

Recent advances in computational methods have made it increasingly feasible to discover and characterize cryptic pockets, particularly through molecular dynamics (MD) simulations^12–18^. For example, Rangari et al. used MD simulations to reveal an extended cryptic pocket in the CB1 receptor located between the orthosteric site and the conserved signaling residue D^2.50^, which subsequently guided the design of a novel ligand^19^. Enhanced sampling techniques have further improved the detection of rare pocket-opening motions, especially those involving slow secondary structure rearrangements. Methods such as Fluctuation Amplification of Specific Traits (FAST)^20,21^, Metadynamics^22,23^, Sampling Water Interfaces through Scaled Hamiltonians (SWISH)^24,25^ have proven particularly effective in this area. In recent studies, the combination of FAST with Markov state models (MSMs), a framework commonly used to map conformational landscapes sampled by MD simulations^26^, successfully identified cryptic pockets in several challenging systems. These included the 5-HT_3A_ receptor, a member of the pentameric ligand-gated ion channel family^27^, and VP35^28^. A follow-up study applying the same strategy to VP35 homologs from Reston and Marburg filoviruses further demonstrated that the open state of this pocket has a biological function, providing an exciting insight into its potential regulatory role^29^. In this study, we found that running FAST simulations followed by seeding additional simulations from evenly dispersed structures discovered by FAST gave reproducible populations of the open state. We refer to this procedure as FAST+seeding in this work to distinguish it from just using FAST simulations without any follow-up simulation to ensure good statistics across the landscape.

While MD simulations are powerful for detecting cryptic pockets, their effectiveness often depends on the specific system, the timescale of the relevant pocket motions, and the choice of simulation parameters^13,25^. The significant computational cost required to achieve adequate sampling can also limit their utility in large-scale or high-throughput studies.

Rapid advances in AI-based methods for structure prediction, such as AlphaFold, Chai-1, and Botz-1, suggest that AI has the potential to accelerate cryptic pocket discovery too^30–34^. For instance, by modifying the multiple sequence alignment (MSA) input, AlphaFold can generate distinct structure variants that may reveal pocket formation directly or serve as starting points for further exploration^35–37^. However, AlphaFold was trained primarily on experimentally determined structures from the Protein Data Bank (PDB)^38^ and large sequence databases, without explicit training to sample alternative conformations. As a result, its ability to capture pocket formation across the global conformational landscape remains limited^35^.

A number of new AI methods have been developed to facilitate broad and efficient exploration of conformational landscapes by using simulations during training. Once such approach, AlphaFlow, repurposes the AlphaFold architecture and incorporates 1.28 million experimentally solved PDB structures^39^. It is further fine-tuned under a custom flow matching framework using the ATLAS dataset, which includes all-atom MD simulations of 1,390 proteins, each performed in triplicate for 100 ns. The resulting model serves as a sequence-conditioned generative framework for sampling protein conformations. This approach achieves faster wall-clock convergence to equilibrium-like properties (e.g., root mean square fluctuation profiles) compared with replicate MD trajectories initiated from a static PDB structure.

Another generative model, BioEmu, was developed using a large and diverse dataset that combines 200 million AlphaFold-predicted structures, over 200 ms of MD simulation data, and approximately 500,000 experimentally measured protein stabilities, integrated through a novel multi-stage training pipeline^40^. Importantly, the BioEmu dataset also includes about 0.5 million mutant sequences, providing a valuable foundation for mutational and stability analyses. The final model accurately predicts equilibrium ensembles and folding free energies, while also capturing dynamic features such as local unfolding events and conformational flexibility in intrinsically disordered proteins.

Beyond generative models, several task-specific predictors like PocketMiner and CryptoBank have been developed for residue-level cryptic pocket detection^41–43^. These models are particularly attractive because they can produce predictions within seconds. One representative structure-based model is PocketMiner, a geometric vector perceptron (GVP)-based graph neural network. It was trained on a curated dataset constructed from 38 proteins containing 39 experimentally validated cryptic pockets^13^, with more than 3,000 independent MD simulations, each lasting at least 40 ns. In total, around 0.9 million residues with simulation-derived labels were used to train the model. This enables PocketMiner to predict whether each residue in a given protein structure is likely to participate in cryptic pocket formation during an MD simulation initiated from that structure. PocketMiner achieved strong performance on the test set (ROC-AUC: 0.87) and is highly computationally efficient. A recently released sequence-based predictor, CryptoBank, fine-tunes a protein language model trained on over 6000 unique sequences derived from 574,000 apo-holo-ligand cryptic alignments. The supervised training dataset used to define cryptic pocket annotations consists of 71 cryptic and 128 non-cryptic cases^44^. The model achieves high precision (PR AUC: 0.8) when query sequences share more than 20% identity with the CryptoBank entries.

Previous work has established that PocketMiner, CryptoBank, and BioEmu can predict where cryptic pockets are likely to form but how well any of these methods predict the equilibrium probability a cryptic pocket is open remains unclear. In this work, we aim to examine the performance of both AI-based and physics-based approaches in capturing the thermodynamics of cryptic pocket opening and the effects of mutations, using VP35 and TEM β-lactamase as model systems (**Figure 1**). These proteins were chosen because their cryptic pockets have been extensively characterized experimentally, with quantitative thermodynamic measurements and detailed studies of mutation-induced conformational changes^9,28,29,45–49^. We specifically evaluate how well each method can distinguish whether mutants lead to pocket opening or closing, quantify the probability of pocket formation, and capture differences among mutants. Through this comparison, we highlight the current limitations of these approaches and underscore the need for further improvements.

**Figure 1.**
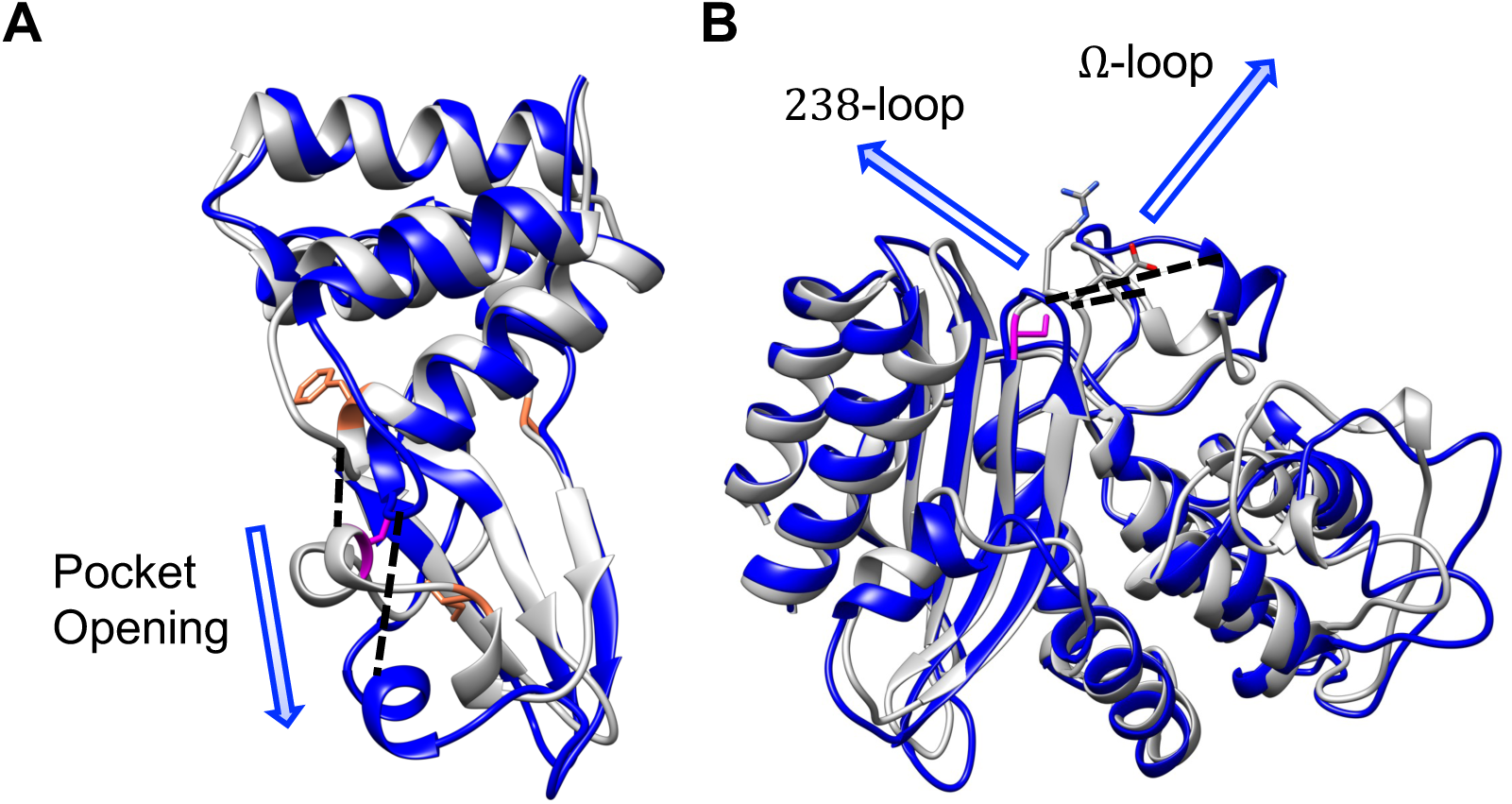
Cryptic pocket opening in VP35 and TEM β-lactamase. **A.** Crystal structure of VP35 (gray; chain A, PDB: 3L25) overlaid with a representative open conformation (blue). The Cα-Cα distance between residues G236 and A306 expands from 5.9 Å (crystal) to 12.1 Å (open), indicated by black dash lines. Residue C307, experimentally labeled and shown to undergo changes in solvent exposure associated with cryptic pocket formation, is colored in magenta in the crystal structure. Mutation sites F239, I303, and A291 are indicated with orange stick representations. **B.** The crystal structure of TEM β-lactamase (gray; PDB: 1JWP) is shown with an open conformation overlaid in blue. The Ω-loop (residue 165-175) and the adjacent 238-loop (residues 237-244) are indicated with arrows. The Cα-Cα distance between E171 and E240 increases from 5.4 Å in the crystal structure to 11.2 Å in the open conformation (black dashed line). Residue S243, mutated to cysteine in thiol labeling assays and associated with increased solvent exposure in the open state, is shown in magenta. Mutation sites E240 and R241 are displayed as sticks.

## Results

### AI-Based and Simulation-Based Methods Successfully Predict Whether Mutants Open or Close Pockets

To evaluate whether computational approaches can reliably detect cryptic pocket formation and predict the effects of mutations in pocket opening, we first focus on VP35 and its mutants, F239A, I303A and A291P (**Figure 1A**). In our previous study^28^, we identified a cryptic pocket in VP35 that forms when a small helix (helix 5) separates from an adjacent four-helix bundle, exposing residue C307. This pocket opening can be captured experimentally using thiol labeling assays, which quantify changes in the solvent accessibility at C307. This study, together with a recent work, demonstrates that mutants F239A and I303A enhance cryptic pocket opening, whereas A291P exhibits a distinct effect by significantly reducing pocket formation^28,50^. These findings provide a clear experimental basis for benchmarking computational predictions across both wild-type (WT) and mutant systems. Additionally, identifying which mutants enhance or suppress pocket opening can help prioritize candidates for further drug discovery or detailed structural studies.

We tested a range of computational methods, including two physics-based simulations (FAST and FAST+seeding MD) and four AI-based models (AlphaFlow, BioEmu, PocketMiner, and CryptoBank). For methods that output structural ensembles (MD and generative AI), we defined open vs. closed states using a Cα–Cα distance between residues G236 and A306, applying a threshold of 1.0 nm. For MD-based simulations, population distributions were calculated from MSMs. For generative AI models, we calculated the percentage of structures exceeding the threshold directly. PocketMiner and CryptoBank predict per-residue opening probabilities rather than full structures, so we compared those scores directly across systems. This unified framework allows a fair comparison of methods with different outputs and sampling strategies (**Figure S1**).

Among the AI-based approaches, AlphaFlow struggles to identify open pockets while BioEmu consistently samples a higher fraction of open conformations across VP35 systems. The predicted open-state populations from BioEmu increase from 10.4% [95% CI: 9.9, 10.8] in WT to 12.6% [95% CI: 12.1, 13.2] in F239A and 18.4% [95% CI: 17.8, 19.0] in I303A, with little change observed for A291P (10.2% [95% CI: 9.6, 10.7]) (**Figure 2**, **Table 1**). These values were estimated from three independent runs initiated from different random seeds, with all other parameters following the default settings in the BioEmu GitHub repository (**Figure S2**). While BioEmu captures some trends, the probabilities it predicts differ substantially from the experimental values and cover a narrower dynamic range (10.4% to 18.4% for BioEmu compared to 0.01-60% from experiments). AlphaFlow, on the other hand, samples fewer than 1% open conformations in all systems. Only two slightly open structures were observed for F239A and one for I303A (**Figure 3A, B**), with similar results across all three replicates (**Figure S3**). Sampling convergence for BioEmu and AlphaFlow was assessed by monitoring how the estimated open-state population varied with the number of sampled conformations (**Figure S4**), indicating the reported populations are well converged.

**Figure 2.**
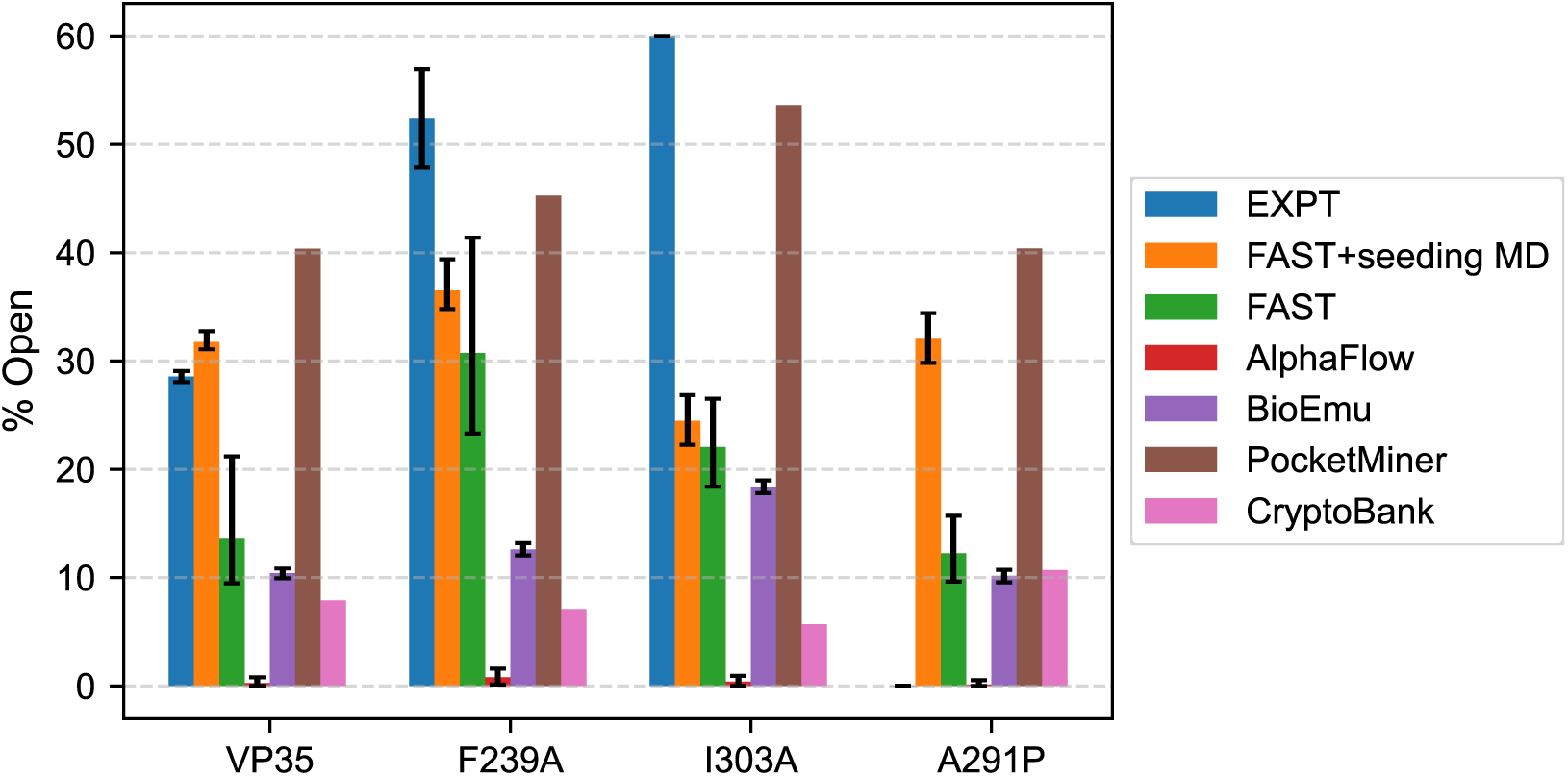
Analysis of cryptic pocket opening in VP35 and its mutants. Computational analysis of cryptic pocket formation in VP35 and three variants (F239A, I303A, and A291P). Open-state populations were quantified using multiple approaches. Experimental values (EXPT) are derived from equilibrium constants measured by thiol labeling assays^28^. Populations from FAST and FAST+seeding MD simulations were calculated by identifying conformations with the Cα-Cα distance between G236 and A306 exceeding 1.0 nm using MSMs. Results from AlphaFlow and BioEmu represent the fraction of sampled conformations meeting the same distance criterion across three replicate simulations per system. Uncertainties for these computational methods are reported as 95% confidence intervals obtained by bootstrap analysis. PocketMiner and CryptoBank predictions indicate the probability of C307 participating in a cryptic pocket.

**Figure 3.**
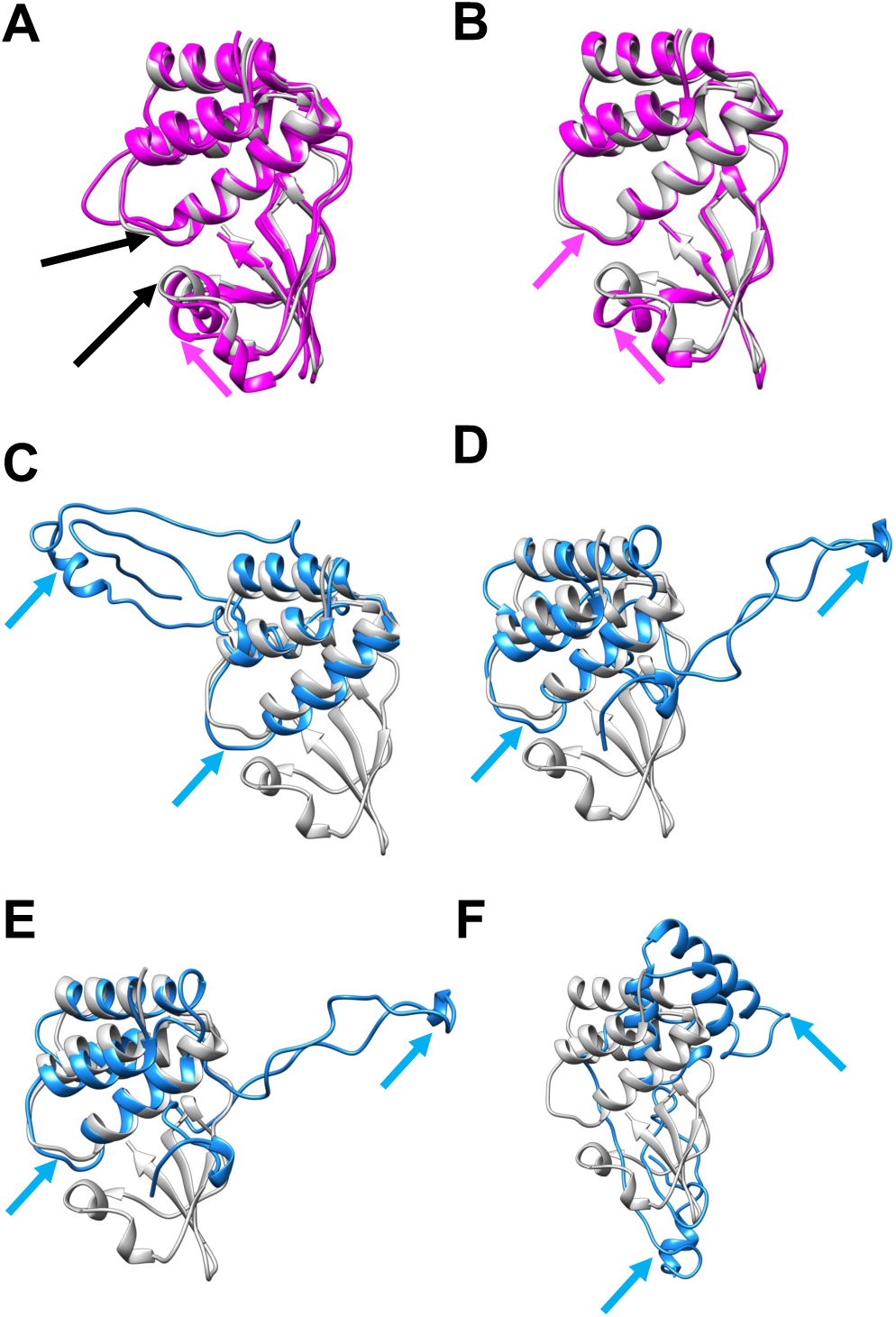
Structural comparison of sampled open conformations with the crystal structure of VP35 highlight that AlphaFlow only captures partial opening while BioEmu sometimes generates implausible structures. Representative open conformations generated by AlphaFlow and BioEmu are overlaid with the VP35 crystal structure (gray) for visual comparison. Residue G236 and A306, which are used to define open versus closed conformations, are indicated by black arrows in panel **A**. The top row shows the only three open conformations sampled by AlphaFlow: two from F239A (magenta, panel **A**) and one from I303A (magenta, in **B**). Magenta arrows highlight the positions of G236 and A306 associate with pocket opening. The bottom rows representative conformations sampled by BioEmu (blue). Panels **C, D, E** and **F** display the structures with the greatest deviation from the conformational ensembles of WT, F239A, I303A, and A291P, respectively. Blue arrows indicate the positions of G236 and A306.

**Table 1.** Numerical values of open-state populations and associated uncertainties for VP35 and its mutants (F239A, I303A, and A291P) derived from experimental measurements and computational methods. Experimental values (EXPT) are inferred from thiol labeling experiments^28^. Simulation-based results from FAST and FAST+seeding MD are calculated using MSMs, with open conformations defined by a Cα-Cα distance between G236 and A306 greater than 1.0 nm. AlphaFlow and BioEmu results report the fraction of conformations meeting this criterion across three replicas. Uncertainties are reported as 95% confidence intervals obtained by bootstrap analysis (values in brackets). PocketMiner and CryptoBank values correspond to predicted probabilities that C307, labeled in experiments, experiences a solvent exposure change associated with cryptic pocket opening.

| VP35 Variant | Cryptic Pocket Open Population (%) |  |  |  |  |  |  |
| --- | --- | --- | --- | --- | --- | --- | --- |
|  | EXPT | FAST+seeding MD | FAST | AlphaFlow | BioEmu | PocketMiner | CryptoBank |
| WT | 28.6 ± 0.5 | 31.8 [31.1, 32.8] | 13.6 [9.5, 21.2] | 0.27 [0.00, 0.80] | 10.4 [9.9, 10.8] | 40.4 | 7.9 |
| F239A | 52.4 ± 4.5 | 36.5 [34.8, 39.4] | 30.7 [23.3, 41.4] | 0.80 [0.13, 1.60] | 12.6 [12.1, 13.2] | 45.3 | 7.1 |
| I303A | > 60.0 | 24.5 [22.3, 26.9] | 22.1 [18.4, 26.5] | 0.40 [0.00, 0.93] | 18.4 [17.8, 19.0] | 53.6 | 5.7 |
| A291P | 0.01 ± 0.02 | 32.0 [29.8, 34.4] | 12.3 [9.6, 15.7] | 0.13 [0.00, 0.54] | 10.2 [9.6, 10.7] | 40.4 | 10.7 |

To test whether substantially increased sampling could improve AlphaFlow’s performance, we increased the number of generated conformations from the 250 used in the original study to 10,000 per system, matching the number generated for BioEmu. Surprisingly, the open-state fraction decreased further to less than 0.01%, despite a slight increase in the absolute number of open structures (**Figure S5**). This result suggests that AlphaFlow struggles to quantitively detect rare pocket-opening events. The computational cost of the two methods also differs significantly. Generating 10,000 samples with AlphaFlow required more than one day per system. In comparison, BioEmu generated the same number of samples in approximately 100 minutes. Using the same computational resources, AlphaFlow required about 70 minutes to produce its original 250-sample run.

These results likely stem from each model’s training strategy and scope. AlphaFlow, which integrates structural prediction and short-timescale dynamics sampling, may be limited by its relatively short simulation lengths and the limited protein diversity in its training set. This could constrain its ability to capture broader or rarer conformational changes. BioEmu incorporates more extensive structural, simulation, and experimental data, allowing it to model equilibrium ensembles more effectively and better reflect conformational flexibility across diverse proteins.

To benchmark physics-based sampling, we performed MD simulations using the FAST adaptive sampling algorithm. FAST accelerates the exploration of targeted conformations by prioritizing trajectory selection based on user-defined features without biasing the underlying potential energy surface. When integrated with MSM analysis, this framework enables efficient sampling of rare conformational states while recovering the true Boltzmann distribution for structured globular proteins^51^. We note, however, that this strategy may be less effective for proteins with substantial intrinsic disorder, such as ApoE, where suitable structural descriptors are difficult to define^52^. Applying this framework to VP35, we utilized FAST to maximize pocket volumes, thereby accelerating the discovery of states with open pockets. Our analysis revealed higher populations of open conformations in F239A and I303A (30.7% [95% CI: 23.3, 41.4] and 22.1% [95% CI: 18.4, 26.5], respectively), along with a moderate decrease in A291P (12.3% [95% CI: 9.6, 15.7]), compared to 13.6% [95% CI: 9.5, 21.2] in WT (**Figure 2**, **Table 1**). So FAST alone gets the direction of change for variants compared to WT but not the absolute probability of pocket opening or the trends between all variants.

To further enhance sampling, we conducted FAST+seeding MD simulations, launching three 40-ns replicas from 1,000 cluster centers per system (120 µs total) (See Methods). Seeding simulations from FAST runs in this manner helps to ensure most of the landscape observed is sampled well, effectively balancing the targeted exploration typical of adaptive sampling. These simulations captured enhanced pocket opening in F239A (36.5% [95% CI: 34.8, 39.4]) and captured WT behavior well (31.8% [95% CI: 31.1, 32.8], compared to the experimental 28.6 ± 0.5%) (**Figure 2**, **Table 1**). A291P displayed a similar level of opening as WT (32.0% [95% CI: 29.8, 34.4]). Interestingly, I303A yielded a notably lower predicted open-state population (24.5% [95% CI: 22.3, 26.9]) relative to the experimental value (> 60.0%). The level of agreement with experiment for WT is encouraging but seeding simulations from adaptive sampling does not correct all the imperfections in adaptive sampling alone.

Finally, we evaluated PocketMiner^13^ and CryptoBank, both of which predict residue-level pocket-opening probabilities from either static structures or the protein sequence, respectively. PocketMiner correctly identified increased opening propensity at F239A and I303A, though the probabilities it predicts are systematically higher than in experiment. It fails to capture the significant reduction in the probability of pocket opening for A291P (**Table 1**). Nonetheless, this performance is encouraging since PocketMiner wasn’t explicitly trained to predict thermodynamics and its training data doesn’t include multiple variants of any proteins. We also tested CryptoBank^44^ by submitting the sequences of each mutant separately. CryptoBank predicted low normalized pocket-opening probability for these variants (WT: 7.9%; F239A: 7.1%; I303A: 5.7%; A291P: 10.7%) and did not reproduce the experimental direction of mutational effects. This discrepancy may reflect the limited inclusion of mutation-specific information in its training data.

### AI-Derived Structural Ensembles Underestimate Pocket Opening While Sometimes Over-Estimating Probabilities of Extremely Extended Structures

Although all the methods that generate structures largely captured the trend of pocket opening from WT to VP35 mutants, the sampled conformational landscapes varied substantially across different ensemble generation approaches. To better understand these differences, we constructed two-dimensional histograms of sampled conformations from each method: FAST+seeding MD, FAST, AlphaFlow, and BioEmu. Each plot maps the inter-residue distance between G236 and A306 (used to define open and closed conformations) against the solvent-accessible surface area (SASA) of residue C307 (**Figure 4**).

**Figure 4.**
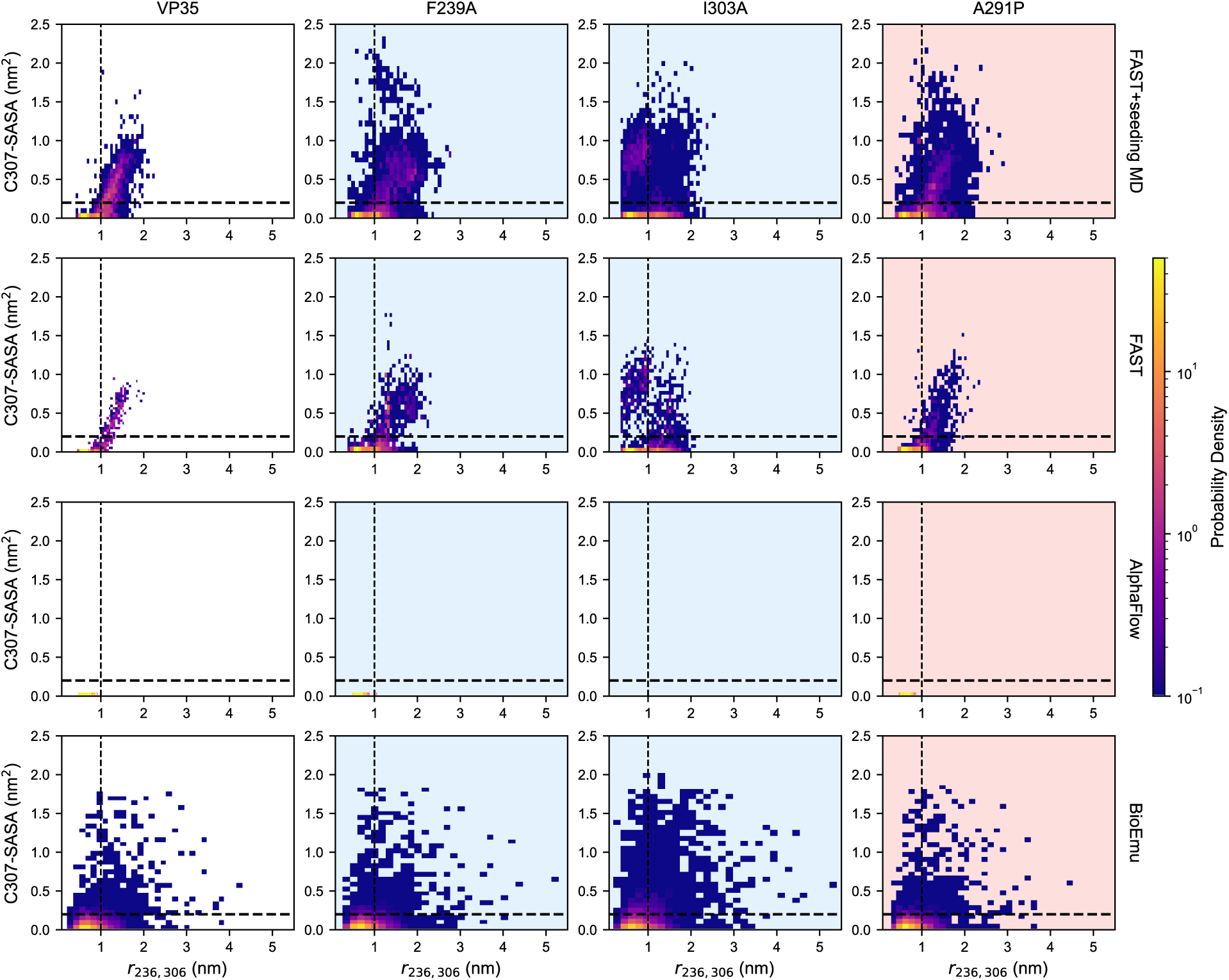
Two-dimensional histograms of sampled conformations generated by each computational method. Each panel shows the inter-residue distance between G236 and A306 (x-axis) plotted against the SASA of residue C307 (y-axis) for VP35 and its three mutants: F239A, I303A, and A291P. Rows correspond to different methods: FAST+seeding MD (top), FAST (second), AlphaFlow (third), and BioEmu (bottom). Background shading highlights whether a mutant opens more or less than WT: light blue indicates variants that open more than WT (F239A and I303A), whereas light pink marks the variants that open less or tend to remain closed (A291P). Logarithmic normalization (LogNorm) is applied to highlight both high- and low-density regions. Dashed lines at x = 1.0 nm and y = 0.2 nm^2^ indicate the distance threshold for pocket opening and (partial) solvent exposure of C307, respectively.

AlphaFlow produced conformations largely similar to the crystal structure, with minimal sampling of open states. Only F239A and I303A exhibit sparse sampling of open states, with just two and one slightly open conformations, respectively (**Figure 3A, B**). WT and A291P display no detectable pocket opening. BioEmu generates broader ensembles across all systems (**Figure 4**). Although the 2D histograms do not reveal dramatic visual differences across systems, I303A shows a notably higher number of conformations with increased SASA at residue C307. This may be attributed to the I303A mutation, which replaces a bulky hydrophobic isoleucine with a smaller alanine. The substitution likely increases the flexibility of the surrounding loop, particularly near the mobile short helix 5, even in closed states where the distance remains below the 1.0 nm threshold.

A subset of BioEmu-generated conformations showed extreme structural deviations that are inconsistent with experiments. Across all systems, 4-15% of BioEmu samples had G236-A306 distances exceeding 3.5 to 6 nm (**Figure S2**), whereas the maximum distance in seeded MD simulations was only 3.0 nm. Inspecting these structures revealed that they have unfolded β-sheets that extend into solvent and are largely devoid of secondary structure (**Figure 3C-F**). The proposed population of these states is inconsistent with ^50^hydrogen-deuterium exchange mass spectrometry (HDX-MS) experiments, which indicate that the population of such extended states is far less than 9.1× 10^-5^%^50^.

Simulation-based methods (i.e., FAST+seeding MD and FAST) also revealed greater conformational flexibility within the closed-state ensemble of I303A (**Figure 4**). This was especially evident in the seeded MD, which sampled a denser and broader range of conformations within the closed state, yet with increased solvent exposure. Such enhanced flexibility and partial exposure may explain the discrepancy between the predicted and experimental open-state populations in I303A (**Table 1**). For F239A and A291P, the FAST+seeding MD results resembled those of the WT, primarily exploring open and solvent-exposed conformations while showing limited sampling closed but exposed states. This suggest that F239A and A291P maintain pocket dynamics more like WT, whereas I303A may undergo alternative or more complex conformational transitions that are not fully captured in our current simulations.

Taken together, all methods distinguished open from closed mutants, but with varying levels of resolution and sampling depth. AI models offer rapid predictions and initial assessments in seconds (PocketMiner or CryptoBank) to hours (AlphaFlow or BioEmu), although they are less effective at resolving detailed structural differences. In contrast, both FAST+seeding MD and FAST sampled the conformational landscapes more thoroughly, capturing the structural features associated with pocket opening. Notably, the FAST simulations, with a total sampling time of 8 µs per system, achieved comparable insights to the 120 µs of FAST+seeding MD simulations, underscoring the efficiency of goal-oriented adaptive sampling.

### Computational Methods Diverge in Capturing Rare Pocket Opening in TEM

To further evaluate the predictive performance of various ensemble generation and pocket detection methods, we tested their ability to detect cryptic pocket opening in TEM variants. TEM is known to contain a Ω-loop cryptic pocket, characterized by motions in the Ω-loop (residue 165-175) and the adjacent 238-loop (residues 237-244) (**Figure 1B**). This pocket exhibits only rare opening in WT (population about 1%), making TEM a valuable test case for assessing each method’s ability to detect subtle cryptic conformational changes.

To quantify pocket opening, we used the Cα–Cα distance between residues E171 and E240 as the opening metric, with a threshold of 1.0 nm to define open versus closed states. Experimental thiol labeling measurements served as ground truth for evaluating model performance^46^. We mainly compared predictions from MD (WT only, conventional trajectories without FAST+seeding), FAST, AlphaFlow, BioEmu, PocketMiner, and CryptoBank (**Figure S6**).

Performance for β-lactamase was generally worse than for VP35. AlphaFlow yielded 0.13% [95% CI: 0.0, 0.53] open conformations, and BioEmu gave 2.2% [95% CI: 2.0, 2.4], compared to the experimental value of 1.1 ± 0.2% (**Figure 5**, **Table 2**). However, AlphaFlow predicted populations less than 1% for all systems in this study, so the congruence with experiments in this case seems more due to the fact that the experimental values happened to fall in this range than that AlpaFold worked better for this system. FAST simulations, however, produced a higher open-state population of 9.9% [95% CI: 7.4, 12.3]. This discrepancy could be due to the underlying force field or to MSM construction not fully correcting for the non-Boltzmann distribution of the raw simulation data. To provide a more direct comparison, we reanalyzed previously reported long-timescale conventional MD simulations (∼90.6 µs) initiated from the crystal structure^47^ (See Methods). Our analysis reproduced the experimental value more closely (0.56% [95% CI: 0.1, 1.2]), supporting the consistency of physics-based sampling when sufficient statistics are available.

**Figure 5.**
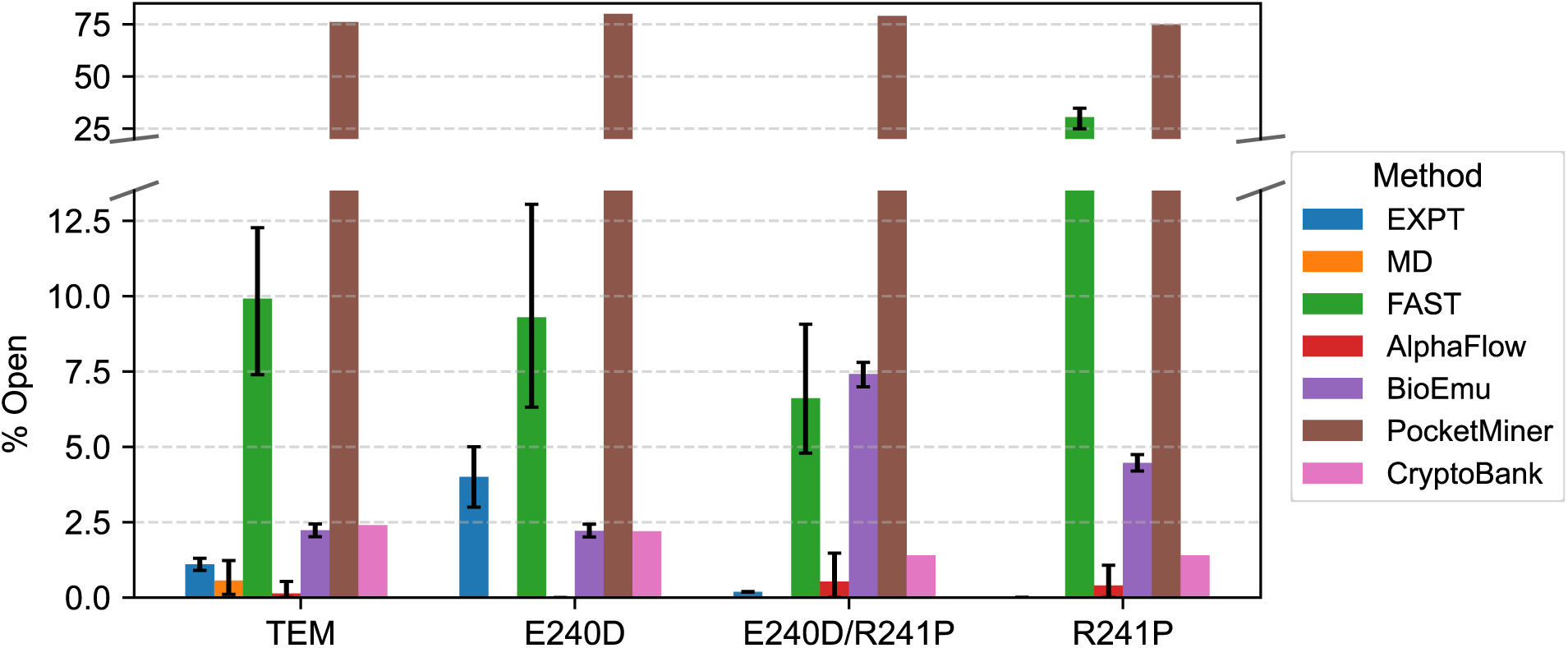
Analysis of cryptic pocket opening in TEM and its mutants. Computational analysis of cryptic pocket formation in TEM and three variants (E240D, E240D/R241P, and R241P). Open-state populations were quantified using multiple approaches. Experimental values (EXPT) are taken from Ref.^46^. Populations from MD and FAST simulations were calculated by identifying conformations with the Cα-Cα distance between E171 and E240 exceeding 1.0 nm using MSMs. Results from AlphaFlow and BioEmu represent the fraction of sampled conformations meeting the same distance criterion across three replicate simulations per system. Uncertainties for these computational methods are reported as 95% confidence intervals obtained by bootstrap analysis. PocketMiner and CryptoBank predictions indicate the probability of S243 participating in a cryptic pocket.

**Table 2.** Numerical values of open-state populations and associated uncertainties for TEM and its mutants (E240D, E240D/R241P, and R241P), derived from both experimental measurements and computational methods. Experimental values (EXPT) are taken from Ref.^46^. Simulation-based results from MD and FAST are calculated using MSMs, with open conformations defined by a Cα-Cα distance between E171 and E240 greater than 1.0 nm. AlphaFlow and BioEmu results report the fraction of conformations meeting this criterion across three replicate simulations. Uncertanties are reported as 95% confidence intervals obtained by bootstrap analysis (values in brackets). PocketMiner and CryptoBank values reflect the predicted probability that residue S243 (mutated to cysteine and labeled in thiol labeling assays) undergoes a change in solvent exposure consistent with cryptic pocket opening.

| TEM Variant | Ω-loop Pocket Open Population (%) | | | | | | |
| --- | --- | --- | --- | --- | --- | --- | --- |
|  | EXPT | MD | FAST | AlphaFlow | BioEmu | PocketMiner | CryptoBank |
| WT | 1.1 ± 0.2 | 0.56 [0.1, 1.2] | 9.9 [7.4, 12.3] | 0.13 [0.0, 0.53] | 2.2 [2.0, 2.4] | 76.3 | 2.4 |
| E240D | 4 ± 1 | - | 9.3 [6.3, 13.1] | 0.0 [0.0, 0.0] | 2.2 [2.0, 2.4] | 80.3 | 2.2 |
| E240D/R241P | 0.19 ± 0.01 | - | 6.6 [4.8, 9.1] | 0.53 [0.0, 1.47] | 7.4 [7.0, 7.8] | 79.0 | 1.4 |
| R241P | 0.002 ± 0.005* | - | 30.5 [24.9, 34.8] | 0.40 [0.0, 1.07] | 4.5 [4.2, 4.7] | 75.0 | 1.4 |

\*Determined to not have an Ω-loop pocket as the pocket populations is less than the unfolded population.

Subtle changes in flexibility proved harder to capture. For example, in the case of E240D, which shortens the side chain from glutamate to aspartate, none of the tested methods effectively detected increased pocket dynamics, including FAST. For the E240P/R241P double mutant, AI-based models incorrectly predicted enhanced opening, while FAST reproduced the experimental trend. For R241P, all methods struggled. FAST significantly overpredicted the open-state population. Among the AI-based predictors, PocketMiner captured the general trend across TEM variants but showed minimal variation between WT and mutants. This suggests that while the model reflects the overall pattern, it may lack the sensitivity to these subtle structural rearrangements. CryptoBank predicted uniformly low normalized pocket-opening probabilities across variants.

To further investigate the sampling behavior of each structural method, we visualized the conformational ensembles generated for TEM and its mutants (**Figure 6**), using the same 2D projection scheme applied in the VP35 analysis. Each plot maps the E171-E240 inter-residue distance (x-axis) against the SASA of S243 (y-axis), a proxy for pocket exposure. Notably, residue S243 was mutated to cysteine and labeled in previous thiol labeling assays^46^. As observed for VP35, AlphaFlow generated a limited conformational ensemble across all three replicates (**Figure S7**). Most sampled structures remained near the closed, crystal-like state, with only a few minor pocket-opening conformations observed among all variants (**Figure 7A-C**). This again suggests that AlphaFlow struggles to capture rare events, particularly when the opening is subtle and infrequent. BioEmu sampled a broader range of conformations but also produces a small number of largely unfolded or extended structures, with E171-E240 distances exceeding ∼4.0 nm (**Figure 7D-F, Figure S8**). These conformations were not found in any MD-based methods. In contrast, FAST explored a more continuous and physically interpretable conformational landscape, effectively targeting the pocket-opening region. These simulations captured intermediate states with partial pocket opening, especially in the double mutant E240P/R241P.

**Figure 6.**
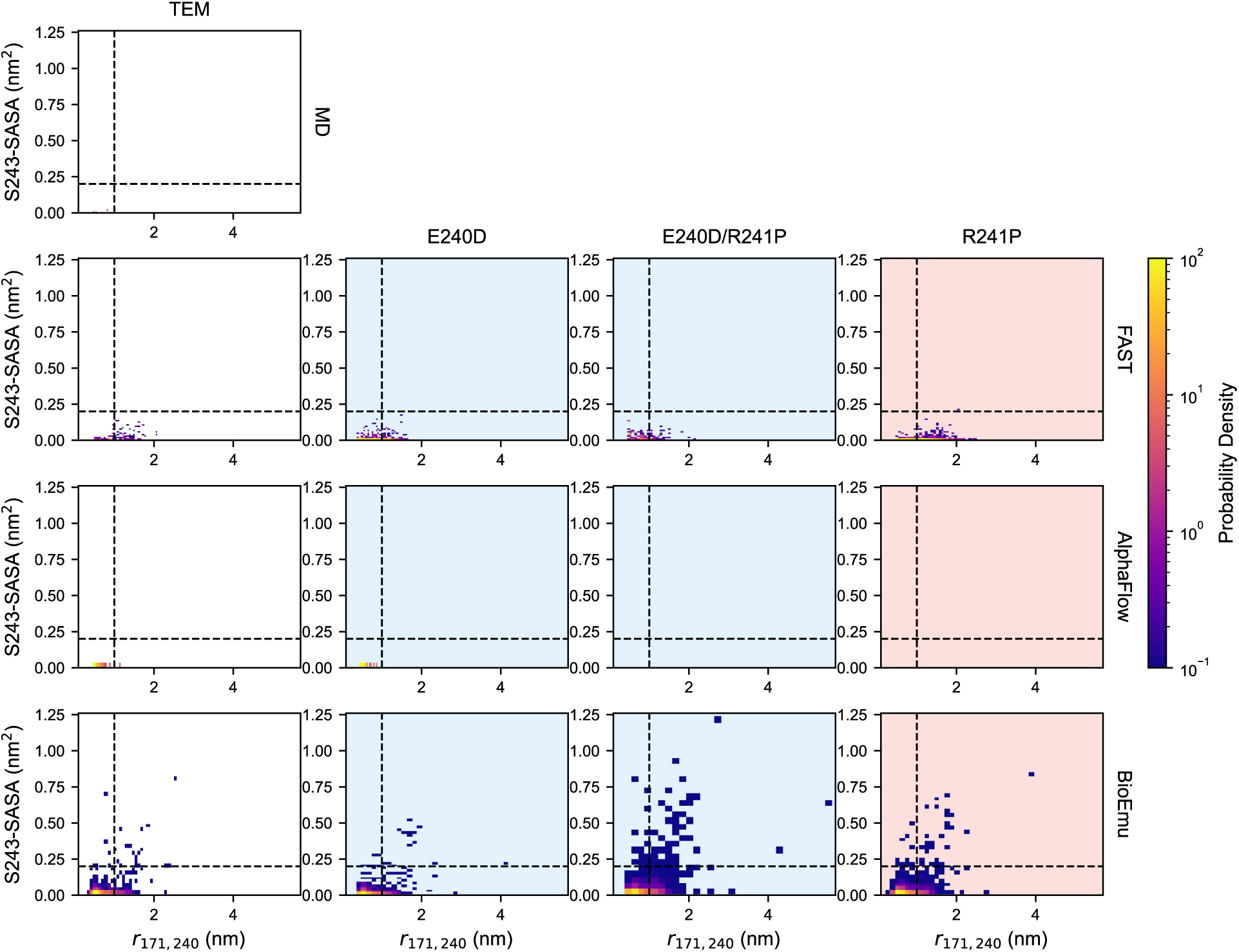
Two-dimensional histograms of sampled conformations from each computational ensemble generation method. Each plot shows the inter-residue distance between residues E171 and E240 (x-axis) versus the SASA of residue S243 (y-axis) for TEM and its three mutants: E240D, E240D/R241P, and R241P. Rows represents different methods: MD (top), FAST (second), AlphaFlow (third), and BioEmu (bottom). Only WT MD simulations initiated from the crystal structure are shown. Background shading highlights whether a mutant opens more or less than WT: light blue indicates variants with increased pocket opening (E240D and E240D/R241P), whereas light pink marks the variant showing no detectable pocket opening (R241P). A logarithmic normalization (LogNorm) is applied to highlight both high- and low-density regions. Dashed lines at x = 1.0 nm and y = 0.2 nm^2^ are included in each subplot to indicate the distance threshold for pocket opening and (partial) solvent exposure of S243, respectively.

**Figure 7.**
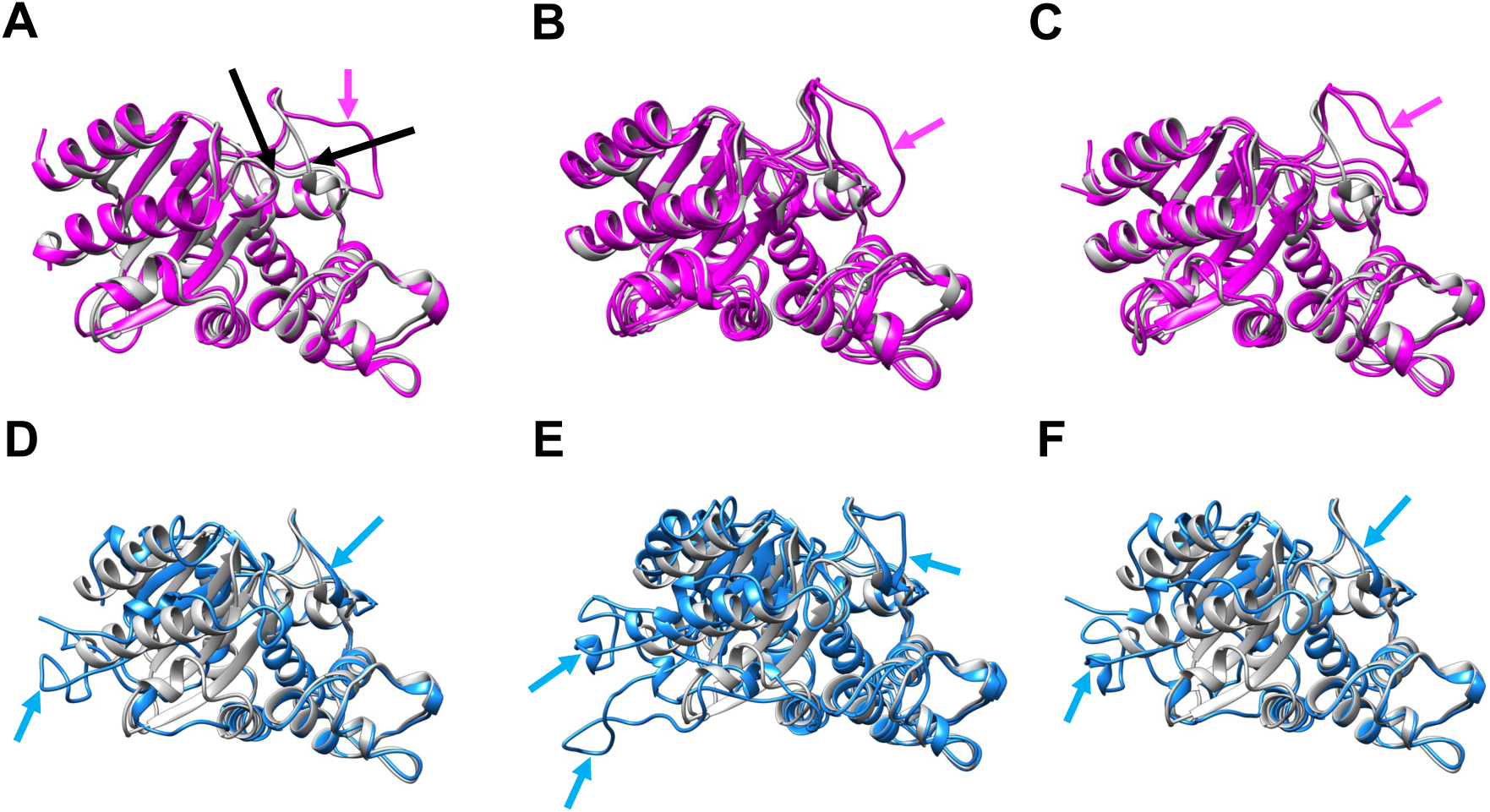
Structural comparison of sampled open conformations with the crystal structure of TEM. Representative open conformations generated by AlphaFlow and BioEmu are overlaid with the TEM crystal structure (gray) for visual comparison. Residue E171 and E240, which define open versus closed conformations, are indicated by black arrows in panel **A**. The top row shows the only six open conformations sampled by AlphaFlow: one from WT (magenta, panel **A**), three from E240D/R241P (magenta, panel **B**), and two from R241P (magenta, panel **C**). Magenta arrows highlight the position of E171, which is associated with Ω-loop pocket opening. The bottom row shows BioEmu-sampled conformations (blue) with inter-residue distances greater than ∼4.0 nm. Panels **D**, **E**, and **F** display these highly open structures from E240D, E240D/R241P, and R241P, respectively. Blue arrows indicate the positions of E171 and E240.

These results highlight the challenges of detecting rare cryptic pockets. While AI-based methods offer computational speed and broad ensemble coverage, they may overlook subtle conformational rearrangements. Conversely, physics-based simulations such as FAST and conventional MD can capture these transitions when sufficient sampling is achieved.

## Discussion

We systematically benchmarked AI-based and physics-based approaches for predicting the thermodynamics of cryptic pocket opening and evaluating mutational-induced effects in two distinct systems: VP35 and TEM β-lactamase. Both proteins have been extensively characterized experimentally, with quantitative thermodynamic measurements on both the wild-type proteins and variants that modulate the probability of pocket opening. The methods tested included goal-oriented adaptive sampling (FAST), FAST+seeding MD, and recent AI models such as AlphaFlow, BioEmu, PocketMiner, and CryptoBank. While prior studies have demonstrated that some of these methods are adept at pocket discovery,^53^ detailed assessments of what these models have learned about thermodynamics has been lacking. Unfortunately, quantitative experimental data on cryptic pocket thermodynamics is only available for a small number of systems. Our focus on two such proteins limits the generalizability of our results but helps give some insight into how current tools can be used (ideally in combination with experiments) and where improvements are needed.

Our results show that all sampling-based methods, as well as PocketMiner, can distinguish between mutants in VP35 known to favor open versus closed pocket opening, though with varying levels of resolution and sampling depth. AI models offered rapid predictions, with outputs generated in seconds (PocketMiner and CryptoBank) to a few hours (AlphaFlow and BioEmu). These tools are valuable for high-throughput initial screening, offering qualitative insights into pocket opening and identifying promising directions for further study. However, BioEmu over populated partially unfolded conformations compared to experimental data. In VP35, FAST simulations achieved comparable insights using just 8 µs of total sampling per system, in contrast to the 120 µs required by FAST+seeding MD, underscoring the efficiency of adaptive sampling. However, in system like TEM, where pocket opening is rare, all methods showed limited ability to fully recapitulate the experimental observations.

Together, our results highlight the complementary strengths of AI- and physics-based approaches. At present, the speed of AI-based methods makes them appealing for triaging large numbers of proteins. MD methods can provide additional insight once one has narrowed down to a smaller set of proteins. While combining these methods can build support for trends, more work is needed to reliably predict the equilibrium probability of pocket opening.

## Methods

### Molecular Simulations

#### System preparation

The protein structures of VP35 (chain A, PDB ID: 3FKE) and TEM (PDB ID: 1JWP) used in this study were obtained from the Protein Data Bank (PDB)^38^. Point mutations, including single and double substitutions, were introduced using PyMOL^54^.

Each protein system was solvated in a rhombic dodecahedral box containing TIP3P water molecules^55^, ensuring a minimum distance of 1.0 nm between any protein atom and the box boundary. System topologies were generated with the Amber03 force field^56^ in GROMACS^57^. To neutralize the system, Na^+^ and Cl^-^ counterions were added to achieve a salt concentration of 0.1 mol/liter. Energy minimization was carried out using the steepest descents algorithm with an initial step size of 0.01 nm and terminized when the maximum force fell below 100 kJ·mol^-1^·nm^-1^ or after 500 steps. Equilibration followed for 0.1 ns using a 2 fs time step. All bonds were constrained using the LINCS algorithm^58^ and long-range electrostatic interactions were treated with the Particle Mesh Ewald (PME) method^59^ with a Fourier spacing of 0.12 nm. A cutoff of 0.9 nm was applied for both Coulomb and van der Waals interactions. The system temperature was maintained at 300 K using the velocity-rescaling thermostat^60^. During both equilibration and production runs, Newton’s equations of motions was integrated with the leap-frog algorithm.

### FAST simulations

We employed the FAST algorithm^20^, which iteratively guides unbiased MD simulations toward conformations that maximize or minimize a chosen order parameter. For VP35 and TEM (both WT and mutant systems), FAST simulations were conducted using pocket volume, calculated by the LIGSITE pocket detection algorithm^61^, as the feature to be maximized. Following the setup described in previous work^28^, each system underwent 10 rounds of adaptive sampling, with 10 parallel trajectories per round, each trajectory running for 80 ns, resulting in a total sampling time of 8 µs per system.

For analysis, the features were defined as the SASA of each residue’s side chain, calculated using the Shrake-Rupley algorithm^62^ implemented in MDTraj^63^, with a drug-sized probe (2.8 Å sphere). Conformations were clustered based on per-residue SASA values using a hybrid k-centers/k-medoids algorithm implemented in Enspara^64^, applying a 2.7 Å distance cutoff for VP35 and 3.0 Å for TEM, followed by five rounds of k-medoids updates. A Markov time of 2 ns was selected based on implied timescales convergence to compute populations with MSMs. The resulting cluster centers were subsequently used to generate the 2D conformational plots shown in the main text; notably, the SASA of C307 presented in these plots was also calculated using the same 2.8 Å probe to ensure consistency.

### FAST+seeding MD simulations

After completing the FAST simulations, conformations were clustered based on Cα RMSD, and 1000 representative structures were used as starting points for additional MD simulations. For each cluster center, three independent simulations were launched, resulting an aggregate simulation time of 120 µs. For the WT of VP35, the simulations were previously performed on the Folding@home distributed computing platform^65,66^ and reported in our earlier study^28^. The VP35 mutant simulations were newly generated on our local high-performance computing cluster, following the same simulation protocol. A corresponding MSM analysis was then carried out for each mutant to extract populations and conformational distributions, using a 3.0 Å distance cutoff and a Markov time of 5 ns (**Figure S9**).

For TEM, we reanalyzed previously reported MD trajectories initiated from the crystal structure^47^ to ensure consistency with our current analysis framework. A Markov time of 25 ns was used for the MSM population calculations (**Figure S10**).

### Experimental Measurements

The experimentally measured results for VP35 and its mutants used in this study were obtained from Ref.^28^. For each system, the open-state population was calculated based on the experimentally determined equilibrium constants describing the exposure of residue C307, using a two-state model.

For TEM, the open-state populations were directly taken from Ref.^47^. Detailed descriptions of the experimental procedures can be found in the respective references.

### AI-Based Models

#### AlphaFlow

Conformations were generated using the AlphaFlow-MD model following the default settings provided in the official GitHub repository. For each system, we first generated 250 samples to match the protocol in the original study, and then increased sampling to 10,000 conformations for testing.

### BioEmu

Conformations were generated using BioEmu with default parameters from the official GitHub repository. To obtain independent replicates, we varied the random seed by modifying batch_size_100. For each system, we generated 10,000 conformations following the protocol described in the reference study.

### PocketMiner

Residue-level pocket-opening probabilities were computed using the provided xtal_predict.py script from the PocketMiner repository, using default settings.

### CryptoBank

Sequence-based predictions were obtained by submitting each mutant sequence individually through the CryptoBank web server.

## Data Availability

The AlphaFlow- and BioEmu-generated data, along with all associated analysis files, have been deposited in the Open Science Framework (OSF) database at https://osf.io/vtx4h. Referenced structures include PDB IDs 3FKE and 1JWP. The starting WT structures used for FAST and MD simulations, as well as the resulting MSM data, are also available in the same directory. Additional FAST and MD simulation trajectories and related files can be provided upon reasonable request.

## Code Availability

FAST, Enspara, and PocketMiner software packages are freely available on GitHub at https://github.com/bowman-lab/fast, https://github.com/bowman-lab/enspara, and https://github.com/Mickdub/gvp.

## Acknowledgements

This work was funded by the National Institutes of Health through NIGMS R35GM152085, and NSF MCB 2218156.

## Competing Interests

GRB consults for VidaVinci and Delphia Therapeutics. The remaining authors declare no competing interests.

## Supporting Information

Distributions of inter-residue distances used to define pocket opening for VP35 (G236–A306) and TEM (E171–E240) across methods; two-dimensional histograms of pocket-opening distance versus residue SASA for VP35 (C307) and TEM (S243) from three independent AlphaFlow and BioEmu replicas; sampling-convergence analysis of predicted open-state populations for AlphaFlow and BioEmu; and MSM implied timescale plots supporting lag-time selection for VP35 mutants and TEM WT.

## Table of Contents

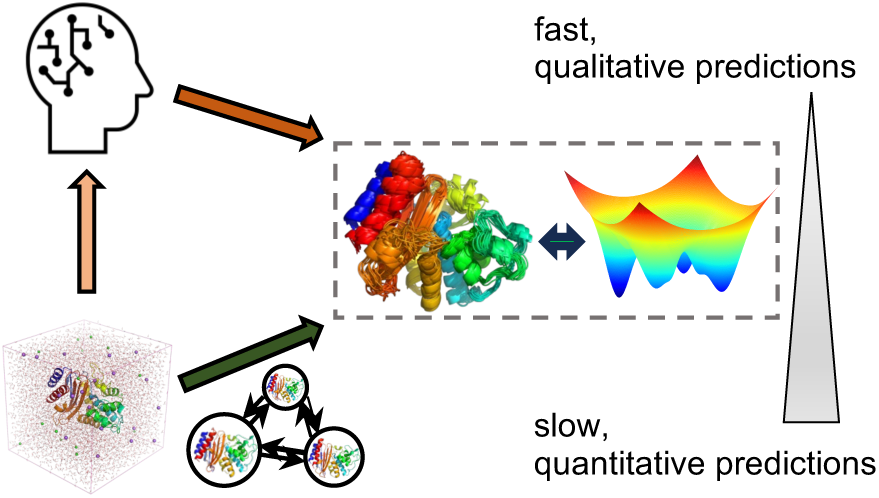

## Notes

### Summary of Updates

Clarified key points and added additional detail

